# TRIM28 regulates Kaiso SUMOylation

**DOI:** 10.1101/2020.12.09.418111

**Authors:** G. Filonova, Y. Lobanova, D Kaplun, S Zhenilo

## Abstract

Tripartite motif protein 28 (TRIM28), a universal mediator of Krüppel-associated box domain zinc fingers (KRAB-ZNFs), is known to regulate DNA methylation of many repetitive elements and several imprinted loci. TRIM28 serves as a scaffold unit that is essential for the formation of stable repressor complexes. In the present study we found that TRIM28 is a binding partner for methyl-DNA binding protein Kaiso. Kaiso is a transcription factor that belongs to the BTB/POZ -zinc finger family. Recent data suggest that deficiency of Kaiso led to reduction of DNA methylation within the imprinting control region of H19/IGF2. Thus, we hypothesized that Kaiso and TRIM28 may cooperate to control methylated genes. We demonstrated that Kaiso interacts with TRIM28 via its two domains: BTB/POZ and three zinc finger domains. When bound to Kaiso’s zinc finger domains TRIM28 weakens their interactions with methylated DNA *in vitro*. Specific association of TRIM28 with BTB/POZ domain causes Kaiso hyperSUMOylation. Altogether our data describe a putative role of TRIM28 as a regulator of Kaiso activity.

## Introduction

DNA methylation is pivotal modification for many processes: vertebrate development, tissue differentiation, cellular reprogramming, X inactivation, gene imprinting. It is crucial for transcriptional regulation of various genes including tumor suppressor genes. DNA methylation most often occurs at the 5’ position of cytosine in CpG context. The presence of methylated cytosines in promoters correlates with a lack of transcription. Removal of DNA methylation in eukaryotic cells results in activation of different repeating elements, reorganization of chromatin accessibility (Cusack et al. 2020). 70% of CpG dinucleotides in the mammalian genome are methylated. Most unmethylated CpG dinucleotides are located in CpG islands (CGIs) and intersected with promoter regions. Methylation of intergenic or located in gene bodies CGIs depends on transcriptional activity of genes or ability to initiate transcription by itself (Jeziorska et al. 2017).

DNA methylation is established and maintained by DNA-methyltransferases 1, 3A and 3B (Goll and Bestor 2005). Depending on the type of sequence: active, poised or inactive promoter, gene body, CpGI, imprinted loci, various repetitive sequences, the mechanism of deposition and maintenance of methylation will be different (Weinberg et al. 2019), (Yagi et al. 2020)) for review (Greenberg and Bourc’his 2019). KRAB zinc finger proteins play a key role in proper DNA methylation and histone mark formation on imprinted loci and various repeating elements (Li et al. 2008) (Alexander et al. 2015). KRAB proteins protect these loci from TET-dependent demethylation (Coluccio et al. 2018). The most investigated member of this family is Zfp57. Zfp57 binds with methylated DNA via zinc fingers and recruited Tripartite motif protein 28 (TRIM28) by KRAB domain. Multicomponent complex is assembled on TRIM28, leading to the di- and trimethylation of H3K9 and DNA hypermethylation. In addition to the role of TRIM28 in the establishment and maintenance of inactive chromatin, it can act as E3 ubiquitin ligase via RING domain located at the N-termini (Doyle et al. 2010). Unlike RING domains in other proteins, the RING domain of TRIM28 can’t dimerize (Stoll et al. 2019). This does not prevent it from participating in ubiquitination and contributes to the fact that it can act as an E3 SUMO ligase (Liang et al. 2011). On the other hand TRIM28 can play an opposite role of epigenetic activator during Th17 cell differentiation (Jiang et al. 2018). The aim of our work was to find new binding partners for TRIM28 that may be involved in maintenance of novel DNA methylation sites. It was shown that another methyl DNA binding protein Kaiso is involved in maintaining DNA methylation of ICR1 in H19/IGF2 imprinted locus and in Oct4 promoter in mouse embryonic fibroblasts (Bohne et al. 2016) (Kaplun et al. 2019). But the mechanism of this maintenance remains unclear. Kaiso belongs to family of protein contains BTB/POZ domain responsible for protein-protein interaction and three C2H2 zinc fingers that can bind methylated CpG and unmethylated CTGCNA sequences, hydroxymethylation prevents binding (A. Prokhortchouk et al. 2001; Daniel and Reynolds 1999; S. V. Zhenilo, Musharova, and Prokhorchuk 2013). Kaiso can attract corepressor complexes NcoR, SMRT (Yoon et al. 2003; Raghav et al. 2012). In this work we showed that Kaiso is a new binding partner for TRIM28. Kaiso interacts with TRIM28 via BTB/POZ domain and zinc fingers. TRIM28 can regulate Kaiso SUMOylation that is important for its transcriptional activity.

## Materials and methods

### Plasmid constructs

Full length Kaiso (1-692), BTB/POZ domain (1-117 amino acids), zinc fingers (494-573 amino acids), spacer (117-494 amino acids) were cloned in pGex-2T.

pKH3-TRIM28 was a gift from Fanxiu Zhu (Addgene plasmid # 45569 ; http://n2t.net/addgene:45569; RRID:Addgene_45569) (Liang et al. 2011). TRIM28 was subcloned to pGex-2T via PCR (for 5’ taagcttctgcaggtcgactctaga 3’ rev 5’ catcgattgaattctcaggggccat 3’).

### Cell culture and transfection

HEK293 was grown in Dulbecco’s modified Eagle medium supplemented with 10% fetal bovine serum, 1% penicillin/streptomycin, and 2 mM L-glutamine. Cells were transfected with Lipofectamine2000 (ThermoFisherScientific) according to the manufacturer’s protocol and were harvested 48 h post-transfection for further analysis. Treatment of cells with MG132 (M8699, Sigma) was performed 30h post-transfection by addition of 0.5, 2, 10 μmol MG132 overnight.

### Protein expression and purification

GST-containing proteins were expressed in E. coli BL21 cells induced by 0.1 mM IPTG for 3 hours at 20 °C in medium with 10 mM ZnSO4. After cell sonication proteins were affinity purified using a Glutathione sepharose 4b or remains on sepharose for pull down (GE Healthcare).

### GST-coprecipitation

GlutathioneSepharose 4B-conjugated BTB/POZ, zinc finger, spacer domains of Kaiso (≈20 μg) were incubated with RIPA total lysates from HEK293 cells transfected with TRIM28-HA during 3 h at 4 °C, washed 4 times with wash buffer (10 mM Tris pH 7.4, 1 mM EDTA, 1 mM EGTA, 150mM NaCl, 1% Triton X-100) and eluted by addition of SDS-loading buffer.

### Antibodies

Following reagents were used in this study: anti-Kaiso polyclonal rabbit antibodies (kindly gifted by Dr. A. Reynolds), anti-HA (H6908, Sigma) anti-HA agarose (A2095, Sigma), anti-actin (ab8227), 6F anti-Kaiso (ab12723, Abcam), anti-KAP1 (ab10483, Abcam).

### EMSA

The binding was carried using Kaiso without BTB/POZ domain (113-692aa) -GST and TRIM28-GST purified from transformed BL21 E.coli using GST-sepharose (Glutathione Sepharose 4B, Cytiva, UK).

Probe was obtained by annealing of methylated oligonucleotides labeled by FAM met-SM-FAM 5’-FAM-GCAGC(meC)G(meC)GCCCAA(meC)GCTGGGAGATC-3’ met-SM 5’-TCCCCAG(meC)GTTGGG(meC)G(meC)GGCTGCGATC-3’(сиквенсы SM пробы оба прайMера).Binding reaction was performed using LightShift EMSA Optimization and Control Kit (20148X) (Thermo Fisher Scientific, USA). DNA-protein complex was loaded to 5% PAAG (0,5X TBE). Resolved complex was detected using Typhon Trio+.

## Results

### Kaiso interacts with TRIM28 via BTB/POZ and zinc finger domains

Kaiso is involved in DNA methylation maintenance on several locus, in repressive chromatin marks establishment ((Bohne et al. 2016; Kaplun et al. 2019; S. Zhenilo et al. 2018)). So, we assumed that Kaiso may be a partner for TRIM28. We checked whether Kaiso can form a complex with TRIM28. HEK293 cells were transfected with Kaiso-GFP and TRIM28 tagged with HA. Immunoprecipitation with antibodies against Kaiso revealed that both proteins immunoprecipitated with each other (fig.1a). To confirm this we performed immunoprecipitation with antibodies against HA. Immunoprecipitation with HA antibodies reaffirmed that Kaiso and TRIM28 formed a complex (fig.1a). Consequently, Kaiso and TRIM28 may exist in one complex or even interact with each other directly.

Next, we analyzed which domain of Kaiso is responsible for complex formation with TRIM28. We obtained GST-sepharose with full length Kaiso, BTB/POZ domain, zinc fingers, spacer region and empty GST as a control. Pull-down was performed with lysates from HEK293 transfected with TRIM28-HA. Western blot of coprecipitated samples confirmed that full length Kaiso interacts with TRIM28 (fig.1b). We detected that BTB/POZ domain and zinc fingers of Kaiso interact with TRIM28, while spacer region and negative control empty GST did not form a complex with TRIM28.

**Figure1.**
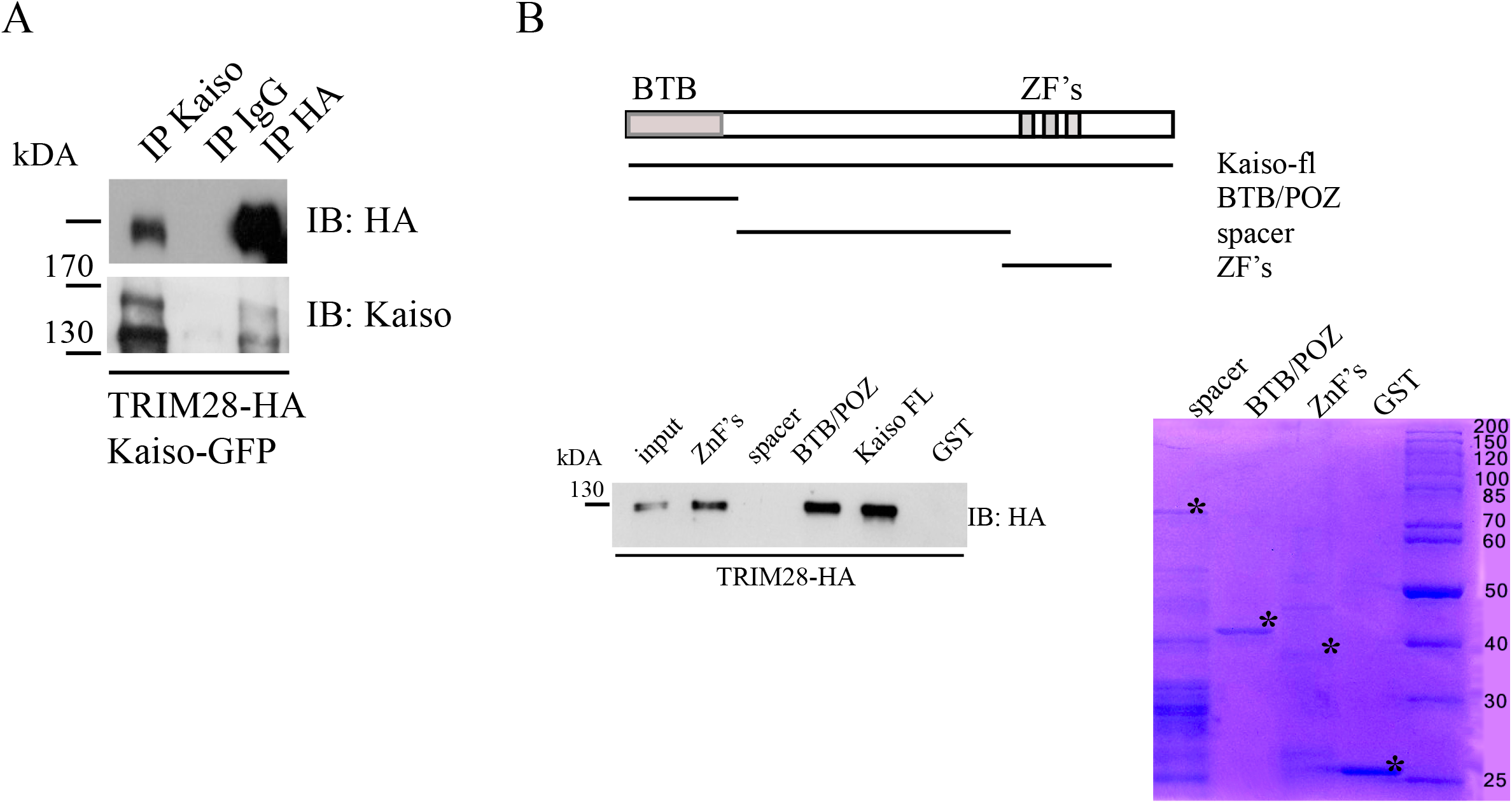
Kaiso forms a complex with TRIM28. a, Kaiso-GFP and TRIM28-HA were transfected to HEK293 cells. Coimmunoprecipitation was performed with Kaiso and HA antibodies. Kaiso is coimmunoprecipitated with TRIM28. b, Scheme of GST conjugated deletion constructs of Kaiso used in GST-pull down. BTB domain and zinc fingers of Kaiso coprecipitated with TRIM28-HA.

**Figure 2.**
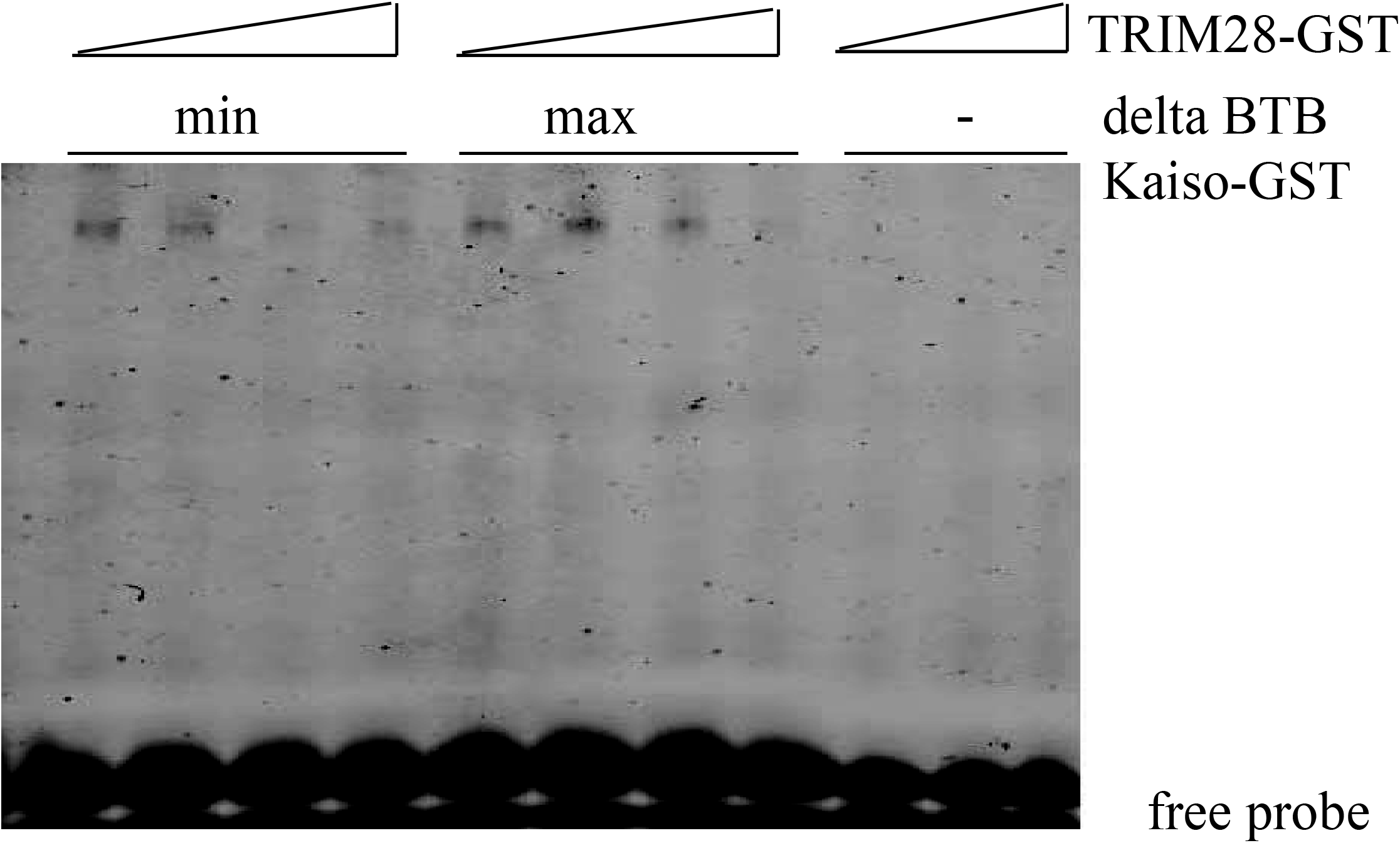
DNA binding activity is affected by TRIM28 protein. High concentration of TRIM28 leads to a decrease in ability of Kaiso binds to DNA. Kaiso without BTB domain conjugated with GST in two concentrations was added to binding reaction with increased concentration of TRIM28-GST.

### DNA binding activity of Kaiso is affected by TRIM28

Zinc fingers of Kaiso can bind DNA and interact with TRIM28. So, it is possible that TRIM28 may influence Kaiso’s ability to interact with DNA. To test this possibility we performed electrophoretic mobility shift assay. Kaiso can interact with TRIM28 via two domains, BTB/POZ domain and zinc fingers. So, we used recombinant Kaiso-GST without BTB/POZ domain. Recombinant Kaiso-GST was incubated with increased concentrations of recombinant TRIM28-GST and labeled by FAM methylated probe. The signal intensity of Kaiso-GST-DNA complex was decreased but not abolished by increased concentration of TRIM28 protein.

### TRIM28 regulates SUMOylation of Kaiso

Further we tested the possibility that TRIM28 is involved in posttranslational modifications of Kaiso. TRIM28 may play the role of E3 SUMO or ubiquitin ligase. We investigated how addition of TRIM28 influences Kaiso SUMOylation. Kaiso is monoSUMOylated by SUMO1. So, we cotransfected Kaiso-GFP with SUMO1-GFP with or without TRIM28-HA. Western blot analysis revealed an increased level of SUMOylated Kaiso in the presence of TRIM28 (fig.3a). Thus, the addition of TRIM28 significantly increased the level of modified Kaiso.

**Figure 3.**
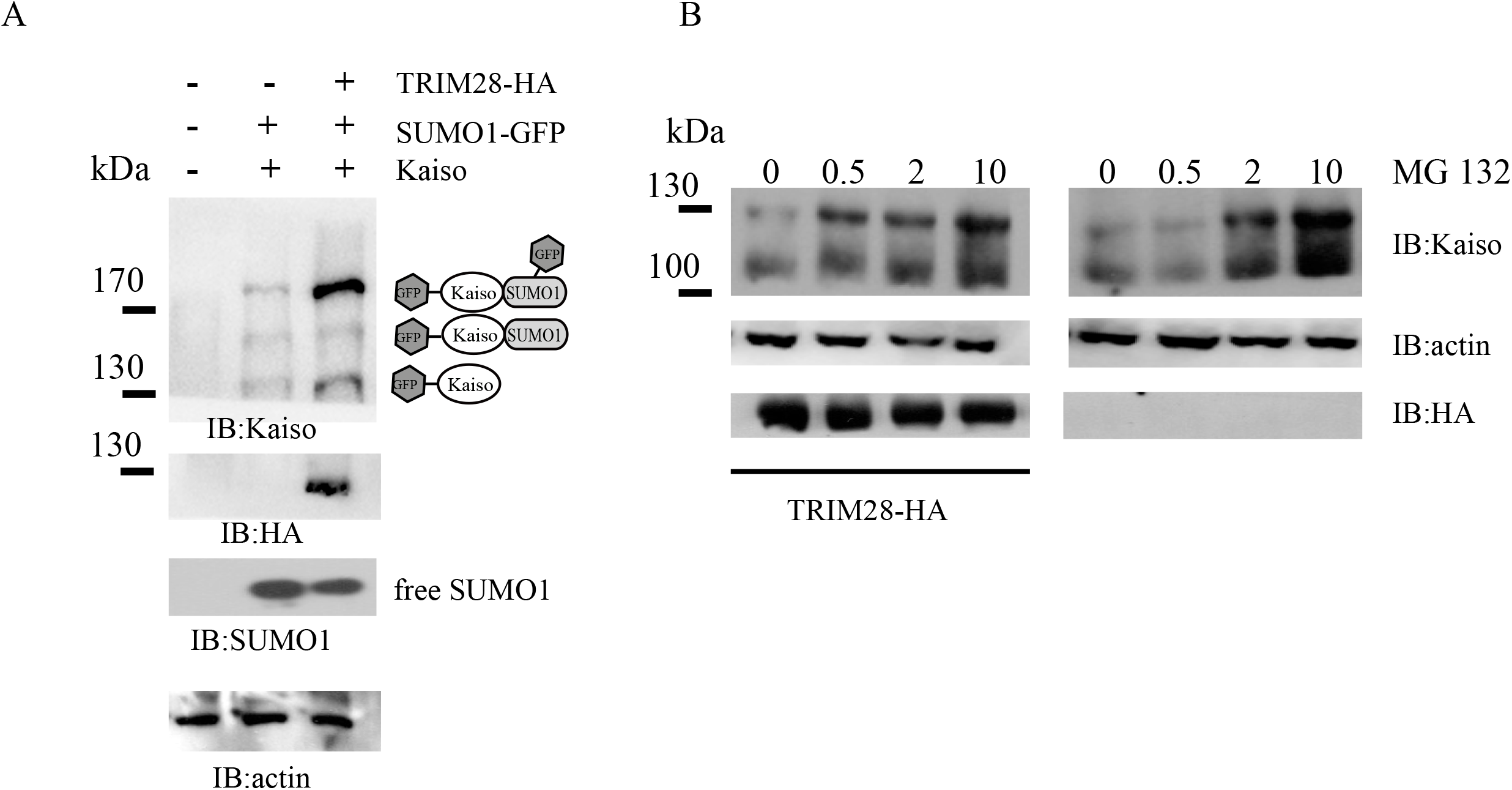
TRIM28 increased Kaiso SUMOylation.a, HEK293 cells were transfected with Kaiso-GFP, SUMO1-GFP and TRIM28-HA. Cells were lysed in the SDS loading buffer. HyperSUMOylation of Kaiso was detected in the presence of TRIM28. b, cells (write panel) and TRIM28-HA (left panel) were treated with increased concentration of MG132 and lysed in the SDS loading buffer. Endogenous Kaiso is stabilized after MG132 treatment regardless of the presence of TRIM28.

Next we analyzed whether Kaiso is a target for ubiquitin machines. Treatment of HEK293 cells transfected with Kaiso-GFP with proteasome inhibitor MG132 lead to stabilisation of Kaiso protein level (fig3b, write panel). Cotransfection with TRIM28 did not change the level of Kaiso protein and did not lead to changes in Kaiso stabilization under MG132 treatment (fig3a, left panel).

## Discussion

In the present study we demonstrated that methyl-DNA binding protein Kaiso can interact with TRIM28 that is involved in transposable elements inactivation and regulation of several imprinted loci. Usually, TRIM28 is attracted to DNA via binding with KRAB methyl-DNA binding proteins. Then TRIM28 acts as a scaffold for DNA methyltransferase, histone methyltransferases and deacetylases. nucleosome remodelling factors that formed a repressive complex on it (David C. Schultz et al. 2002; D. C. Schultz 2001)). Knockout of TRIM28 leads to embryonic lethality ((Shibata et al. 2011)) in accordance with its role in imprinting regulation. Knockout of KRAB protein Zfp57 main partner of TRIM28, is similar to TRIM28 deficiency (Li et al. 2008). So, TRIM28 and Znf57 are closely related and important for mammalian development. TRIM28 is highly conservative between species and can be found in Xenopus laevis and Danio rerio. However, Znf57 or other KRAB proteins that can interact with TRIM28 are absent in Xenopus laevis and Danio rerio. Nothing is known about the necessity of TRIM28 for frog or fish development. This raises a question about the possibility of the existence of another class of proteins in Xenopus that can bind methylated DNA and attract TRIM28 to form repressive chromatin. We hypothesized that this may be a protein Kaiso. Kaiso is essential for Xenopus and fish development, its deficiency leads to premature zygotic gene expression occurring before the mid-blastula transition (MBT) resulted in developmental arrest and apoptosis (Ruzov et al. 2004). This phenotype is similar to observed in DNA methyltransferase xDNMT1 deficiency embryos with hypomethylated genome (Stancheva, Hensey, and Meehan 2001). In mammals the role of Kaiso is not so obvious. Deficiency of Kaiso mainly resulted in behavioral changes (Korostina and Kulikov 2015). Kaiso interacts with the same imprinted methylated loci as Znf57, but its deficiency leads to slightly decreased methylation of this region without changes in imprinted genes transcription and reactivation (Anna Prokhortchouk et al. 2006). Recently we found that Kaiso knockout leads to partial demethylation of Oct4 promoter region in mouse embryonic fibroblasts (Kaplun et al. 2019). Moreover deSUMOylation of Kaiso may lead to heterochromatin formation at TRIM25 promoter (S. Zhenilo et al. 2018).

Here we show that TRIM28 is immunoprecipitated with Kaiso in human HEK293 cells and may play different roles for regulation of Kaiso activity. On one hand, TRIM28 can interact with the BTB/POZ domain of Kaiso. This interaction may have an impact on Kaiso SUMOylation, since Kaiso is modified by SUMO on 42K in BTB/POZ domain. SUMOylation reverted Kaiso from repressor to activator (Zhenilo et al. 2018). On the other hand, this interaction may be important for complex formation. Further research is needed to determine if this complex will be activating or repressive. Also TRIM28 interacts with the zinc finger domain of Kaiso. The zinc finger domain of Kaiso is responsible for DNA binding and is involved in interaction with proteins. So, binding of p120 catenin to the zinc finger domain of Kaiso abolished the complex formation of Kaiso with DNA (Zhigalova et al. 2010). TRIM28 decreased the DNA binding ability of Kaiso but did not abolish it. Consequently, TRIM28 may play different roles for regulation of Kaiso activity.

## Conclusion

Kaiso interacts with TRIM28 via BTB/POZ domain or zinc fingers. TRIM28 is involved in Kaiso SUMOylation. TRIM28 decreases the binding of Kaiso with DNA.

## Acknowledgments

Sanger sequencing was carried out on the equipment of the Center of Collective Use “Bioengineering” of the Federal Research Center of Biotechnology RAS

